# Differential activation states of direct pathway striatal output neurons associated with the development of L-DOPA-induced dyskinesia

**DOI:** 10.1101/2023.11.03.565591

**Authors:** David A. Figge, Henrique de Amaral Oliveira, Jack Crim, Rita M. Cowell, David G. Standaert, Karen L. Eskow Jaunarajs

## Abstract

L-DOPA-induced dyskinesia (LID) is a debilitating motor side effect arising from chronic dopamine replacement therapy with L-DOPA for the treatment of Parkinson disease (PD). The emergence of LID is linked to heightened sensitivity of striatal dopaminergic signaling and driven by abnormal fluctuations in synaptic dopamine levels following each administration of L-DOPA. This maladaptive plasticity narrows the therapeutic window for L-DOPA treatment, even as the progressive worsening of PD symptoms demands escalating doses. The heterogeneous composition of the striatum, including diverse subpopulations of medium spiny output neurons (MSNs), interneurons, and supporting cells, has complicated the precise identification of the cell(s) underlying LID development and persistence. To elucidate the cellular and molecular mechanisms of LID, we used single nucleus RNA-sequencing (snRNA-seq) to establish a comprehensive striatal transcriptional profile during the development and maintenance of LID in an animal model. Hemiparkinsonian mice were treated with vehicle or L-DOPA for progressive durations (1, 5, or 10 d) and nuclei from the striata were processed for snRNA-seq. Our analysis found that a limited population of dopamine D1 receptor-expressing MSNs (D1-MSNs), arising from both the patch and matrix compartments, formed three subclusters in response to L-DOPA treatment that expressed cellular markers of activation. These activated D1-MSN subpopulations display many of the transcriptional changes previously associated with LID; however, the prevalence and transcriptional behavior of activated D1-MSNs was differentially influenced by the extent of L-DOPA experience. The differentially expressed genes found in these D1-MSNs indicated that acute L-DOPA induced upregulation of multiple plasticity-related transcription factors and regulators of MAPK signaling, while repeated L-DOPA exposure induced numerous genes associated with synaptic remodeling, learning and memory, and transforming growth factor-ß (TGFß) signaling. Notably, repeated L-DOPA led to a sensitization in the expression of *Inhba,* a member of the activin/TGFß superfamily, in activated D1-MSNs. We tested pharmacological inhibition of its receptor, ALK4, and found that it impaired LID development. Collectively, these data suggest that distinct subsets of D1-MSNs become differentially responsive to L-DOPA due to the aberrant induction of the molecular mechanisms necessary for neuronal entrainment, similar to those processes underlying hippocampal learning and memory formation. Our data further suggests that activin/TGFß signaling may play an essential role in LID development in this subpopulation of D1-MSNs.

## Introduction

Nigrostriatal dopamine (DA) neuron loss is the cardinal hallmark of Parkinson disease (PD) and is most commonly treated with levodopa (L-DOPA) therapy. While initially beneficial, L-DOPA treatment frequently results in the progressive development of abnormal involuntary movements, known as L-DOPA-induced dyskinesia (LID). At least 40% of patients treated with L-DOPA will experience LID after 5 years of therapy (Ahlskog & Muenter, 2001). LID is believed to stem from an imbalance in the output of striatal medium spiny neurons (MSNs) from the kinetogenic direct (striatonigral) pathway at the expense of the inhibitory indirect (striatopallidal) pathway (Castela et al., 2023; Cenci et al., 2018). L-DOPA administration causes the activation of dopamine D1 receptor-expressing MSNs (D1-MSNs) within the direct pathway to drive LID expression. The biochemical and synaptic sensitization are most prominently found in the D1- MSNs following L-DOPA exposure, while D2-MSNs show minimal changes in function during LID development. Specific manipulation of either pathway can affect LID expression; however, only stimulation of D1-MSNs can independently induce dyskinesia in the absence of L-DOPA (Alcacer et al., 2017; Castela et al., 2023).

In addition to the functional differences of MSNs based upon their dopamine receptor expression, the striatum is somatotopically organized with the centrolateral striatal regions associated with sensorimotor inputs, while the medial regions play a more important role in associative-and goal-oriented behaviors (Burton et al., 2015). Furthermore, the striatum also contains compartments with unique inputs and outputs known as the patch (or “striosome”) and matrix (Brimblecombe & Cragg, 2017). However, due to limitations in isolating specific neuronal populations, the differential roles that these subareas may play and their respective contributions to the precise microcircuitry underlying LID remains unclear.

In response to chronic L-DOPA, D1-MSNs undergo numerous physiological and biochemical changes. Following the induction of LID in animal models, D1-MSN corticostriatal synapses exhibit long-term potentiation (LTP) that has been shown to be uniquely resistant to synaptic depotentiation (Iravani et al., 2012; Thiele et al., 2014). Underlying these synaptic changes is a persistent biochemical sensitization in MAPK signaling following D1 receptor stimulation, causing increased ERK/CREB phosphorylation following L-DOPA administration (Cerovic et al., 2015; Mariani et al., 2019; Pavon et al., 2006; Santini et al., 2009; Santini et al., 2012; Westin et al., 2007). Moreover, the altered dopaminergic signaling leads to long-term enhancements in the evoked transcription of several immediate-early genes (IEGs), including *Zif268*, *Arc*, *Fos*, and *FosB,* exclusively in D1-MSNs following L-DOPA treatment (Bastide et al., 2014; Carta et al., 2008; Fieblinger et al., 2018; Girasole et al., 2018; Westin et al., 2007). This biochemical and transcriptional sensitization is crucial for the manifestation of abnormal behaviors, and inhibition of this sensitization can block LID expression in animal models (Darmopil et al., 2009; Santini et al., 2009). Although enhanced signaling in D1-MSNs has been implicated in LID, a clear understanding of the transcriptional response to L-DOPA across the development of dyskinesiais lacking. In addition, the response of the MSN subpopulations, interneuronal, and non-neuronal populations remains unknown. Thus, a comprehensive understanding of the cellular landscape and transcriptional network affected by L-DOPA exposure remains necessary to fully understand the underlying pathophysiological mechanisms.

Although bulk RNA-seq techniques have identified numerous changes in gene expression following LID development (Dyavar et al., 2020; Smith et al., 2016; Sodersten et al., 2014), due to the cellular complexity of the striatum (Gokce et al., 2016; Martin et al., 2019; Stanley et al., 2020), the particular cells specifically involved remain unknown. Recent advancements in single-nuclei (sn) RNA-seq technologies have facilitated the exploration of complex tissues with significant cellular heterogeneity, enabling the generation of cell type-specific transcriptional profiles during disease modeling and drug exposure. Using snRNA-seq, we studied the kinetics of the striatal transcriptional response during LID development in the 6-hydroxydopamine (6-OHDA) mouse model of PD. Our unbiased approach provides insights into cellular subtypes and gene modules involved in LID entrainment and stable expression, highlighting a key role for specific D1-MSN subpopulations.

## Methods

### Mice

Male C57Bl/6J mice (8 weeks of age) purchased from Jackson Labs were utilized. Mice were group-housed with a 12 h light/dark cycle. Food and water were provided *ad libitum*. All studies were approved and complied with the University of Alabama at Birmingham Institutional Animal Care and Use Committee guidelines and protocols.

#### Experiment 1. Cell type-specific transcriptional profile of LID development

### 6-hydroxydopamine lesion surgeries

To induce hemiparkinsonism, mice were subjected to a unilateral infusion of 6-hydroxydopamine (6-OHDA) into the medial forebrain bundle to deplete striatal dopamine, utilizing standard stereotaxic survival surgery techniques as previously described (Thiele et al., 2012). Mice were treated with desipramine HCl (25 mg/kg, ip) 15 min prior to surgery to protect noradrenergic neurons. 6-OHDA HBr (0.2 uL in 0.2% ascorbic acid and 0.9% saline; 3 ug/uL) was injected with a Neuros 33 gauge 10 uL syringe (Hamilton Apparatus) at the following coordinates: AP,-1.2; L,-1.1; V,-5.0 mm. Mice received buprenorphine analgesic and Lactated Ringer’s solution (sc) to prevent dehydration, and were allowed to recover on a heating pad for up to 8 h post-surgery. Any mice that expressed distress or dehydration during the 7 d recovery period were treated with additional fluids and/or additional heating pad time.

### Vehicle-induced rotations

Two weeks post-surgery, mice were individually placed into clear acrylic cylinders. After 30 min of habituation, mice received an injection of saline (0.2 mL, sc) and were immediately returned to the cylinder. For 5 min post-injection, ipsilateral and contralateral rotations were counted, with a tendency toward ipsilateral rotations considered indicative of hemiparkinsonism. Mice were ranked based on the number of ipsilateral – contralateral rotations and separated into equal groups: Vehicle-treated or L-DOPA-treated for 1 (Acute), 5 (Subchronic), or 10 (Chronic) d.

### Abnormal Involuntary Movements Scale (AIMs) assessment of LID

Three weeks post-surgery, mice commenced 10 d of treatment with either vehicle (0.9% sodium chloride + 0.1% ascorbic acid, Sigma, USA) or L-DOPA methyl ester hydrochloride (4 mg/kg + 15 mg/kg benserazide hydrochloride, Sigma, USA), as indicated by Fig 1A. LID severity was assessed with the AIMs (Lundblad et al., 2002; Lundblad et al., 2005) for 3 h on Days 1 and 5, and 1 h on Day 10, prior to harvesting of striata. In brief, mice were individually placed into the same clear acrylic cylinders utilized for vehicle-induced rotations. After 30 min of habituation, mice received an injection of vehicle or L-DOPA and were returned to the cylinder. Axial, limb, and orolingual (ALO) AIMs were separately quantified after observation for 1 min every 20 min on a scale of 0- 4 (0=absent, 1=present for <30 s, 2=present for >30 s but <1 min, 3=present for 1 min, but interruptible by a tap on the cylinder, 4=present for 1 min, uninterruptable by a tap on the cylinder). Contralateral rotations were also counted, as a traditional measure of differential dopamine sensitivity (Norman et al., 1990).

**Figure 1.**
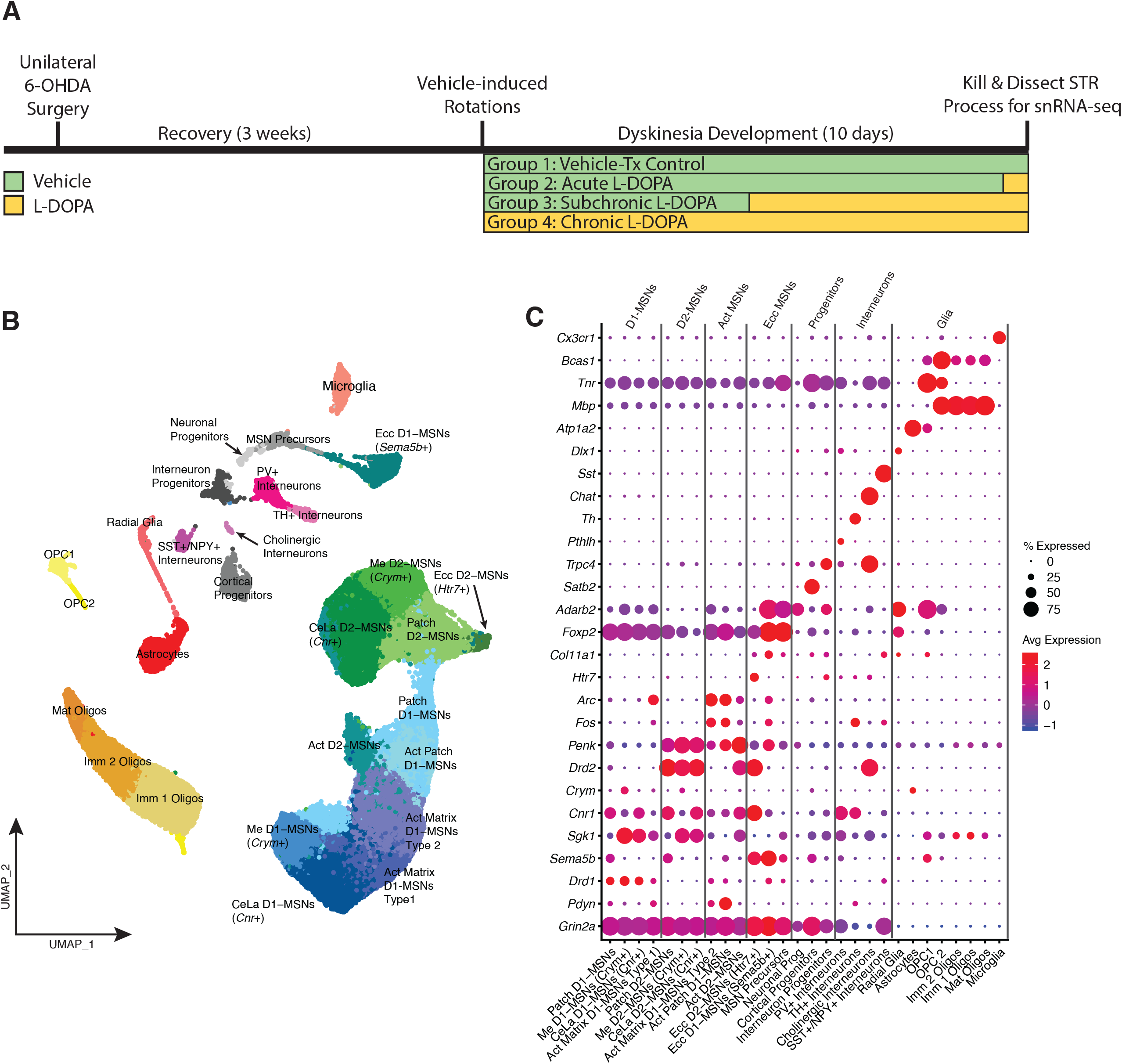
Clustering analysis reveals all major striatal cell subtypes. (A) Time course of experiment and graphical description of experimental groups for snRNA-seq experiments. (B) Integrated UMAP of striatal cellular subclustering across all experimental groups (>97,000 nuclei). (C) Dot plot examining % of nuclei expressing cluster-defining genes (size of dot) and their average expression (color of dot) within each cluster.

### Nuclei preparation and snRNA-seq

One h after the last vehicle or L-DOPA injection, mice were anesthetized with isoflurane and decapitated. The hindbrain was removed and placed into cold 4% paraformaldehyde in 0.1 M phosphate buffered saline for fixation and consequent quantification of dopamine neuron depletion in the substantia nigra. Left (lesioned) and right (intact) striata was removed by microdissection and pooled with 1-3 other mice that were in the same treatment-duration group. Nuclei isolation was as previously published (Savell et al., 2020). The tissue was chopped orthogonally and lysed for 15 min in 15 mL of lysis buffer containing 10 mM Tris-HCl, 10 mM NaCl, 3 mM MgCl2, and 0.1 % Ipegal in nuclease-free water. The reaction was quenched with 5 mL of Hibernate-A (Thermo-Fisher) containing B27, Glutamax (Life Technologies), and 0.2U/uL RNAse inhibitor (Lucigen). The tissue was then triturated 15 times through a fire-polished Pasteur pipette and then passed through a 40 uM filter to remove cell debris and large chunks. The filtered solution was centrifuged at 500 rcf for 10 min, the supernatant removed, and pelleted nuclei washed with 10 mL 1X phosphate buffered saline with 1.0% bovine serum albumin and 0.2U/uL RNase inhibitor. This was centrifuged at 500 rcf for 5 min and the nuclei pellet resuspended in 300 uL of the wash buffer. Nuclei were then tagged with propidium iodide and purified using flow-assisted cell sorting (FACS; BD FACS Aria II, 70 uM nozzle, BD Biosciences) and brought to a concentration of 1000 nuclei/uL. An average total of 10000 nuclei/sample were loaded into each well of the Chromium Single Cell B Chip (10X Genomics). Single nuclei libraries were constructed according to the instructions provided in the Chromium Single Cell 3’ Library Construction Kit (10X Genomics). Nuclei were sequenced on an Illumina Novaseq with a minimum depth of 15000 reads/nuclei. Sequencing files were processed, mapped to mm10, and count matrices were extracted using the Cell Ranger Single Cell Software (v 3.1.0).

### snRNA-seq analyses

All analyses were conducted in R using Seurat (v3.1) as previously published (Schonhoff et al., 2023). Briefly, all data underwent quality control filtering any genes expressed in fewer than 3 cells and excluding cells with <300 unique transcripts or more than 3% mitochondrial genes. The datasets were then integrated based upon the 2000 most variable genes. The data was normalized and the top 20 principal components were used for UMAP dimensional reduction. Clustering was performed following identification of nearest neighbors at a resolution of 0.7. Clusters predominantly composed of either cells with low RNA content or doublet cells were removed and the datasets were re-clustered following dimensional reduction. Marker genes for each cluster were determined using the FindAllMarkers function of Seurat with a minimum Log2 fold change threshold of +/−0.25 with the Wilcoxon ranked-sum test. For the direct comparisons between clusters, we used FindMarkers with similar thresholds. Filtered and merged datasets were imported into Monocle3 (Trapnell et al., 2014). Module analyses were conducted using standard Monocle commands. GO analyses were completed using Metascape (Zhou et al., 2019) and/or PantherDB (Thomas et al., 2022). The dataset was compared to two additional datasets to 1) indicate D1-MSN cell-subtype specificity of prior D1-MSN RNA-seq data in a model of LID (Heiman et al., 2014) and 2) compare the transcriptional profile of D1- MSNs differentially activated by L-DOPA treatment to hippocampal engram cells in a model of learning and memory (Marco et al., 2020). Data from the Marco et al., 2020 dataset were reanalyzed to generate DEGs from nRNA-seq data from initial exposure compared to reactivation as described above. Transcription factor enrichment was determined using LISA Cistrome database TR ChIP-seq combined model (Qin et al., 2020) on the top 200 DEGs (based on LogFC) obtained from pseudobulk-seq analysis of D1-MSN subpopulations.

### Determination of dopamine neuron-depletion

Fixed hindbrain tissue from each mouse was cryoprotected with 30% sucrose in 0.1 M PBS, frozen on dry ice and sliced on a freezing sliding microtome at 40 uM thickness. Every sixth section containing the substantia nigra was collected for immunohistochemistry of tyrosine hydroxylase (TH)+ dopamine neurons. Free-floating sections were washed and exposed to 1% normal donkey serum in Tris buffered saline (TBS) and 0.1% Triton-100. Utilizing a fluorescence-based strategy, slices were exposed to donkey anti-goat TH primary antibody followed by a LiCor Odyssey 700 secondary antibody. Slices from each animal were then mounted on coated glass slides and placed face-down on the scanner surface of a LiCor Odyssey DLx. Slides were scanned at highest quality and resulting images analyzed using ImageStudio Lite freeware. Regions of interest were drawn around contralateral and ipsilateral substantia nigra and the signal/area quantified for each slice. Averages of at least 4 slices were used to establish a % of intact side by the following: average lesioned signal/ average intact signal x 100.

#### Experiment 2. Independent verification of top gene upregulated in chronic versus acute L-DOPA treated D1-MSNs

### Fluorescence in situ hybridization (FISH)

We utilized RNAScope Multiplex Fluorescent Assay (ACD Biosciences, Newark, CA, USA) in a separate group of 6-OHDA-lesioned mice to verify the enhanced expression of *Inhba* in D1-MSNs that also expressed Fos in response to chronic compared to acute L-DOPA treatment. Mice were treated with Vehicle or L-DOPA (4 mg/kg + benserazide, 15 mg/kg, sc) for 1 or 10 d and killed 1 h after injection. AIMs were assessed on days 1 and 5 for 2-3 h and for 1 h on day 10. Brains were removed and flash frozen on dry ice. Sections (20 micron) were obtained using a cryostat, collected on slides, and refrozen immediately. Tissue was fixed with 4% paraformaldehyde followed by ethanol dehydration and treatment with hydrogen peroxide and protease III (ACD). Tissue was then incubated according to prescribed protocol (ACD) with probes targeted for *Inhba, Fos* and *Drd1.* Slides were coverslipped with Prolong Gold Antifade Mounting Media with DAPI (Thermo Fisher Scientific) and images captured at 20X objective with a Nikon confocal microscope (n=4 animals/treatment, 3 sections/animal). All settings were held constant across all groups for a given experiment. Images were analyzed using QuPath (Courtney et al., 2021). Cell detection was performed using DAPI signal following optimized parameters of QuPath defaults. Next, we performed cell classification based on fluorescent intensity thresholds of each of the selected channels: *Inhba* (fluorescein), *Fos* (Cy3), *Drd1* (Cy5). We applied the “object classification function” and created a single measurement classifier for each channel. We labeled cells based on the presence of fluorescence, considering the intensity threshold, for each of the applied channels (*Inhba*, *Fos* and *Drd1*). After this last process, the data was exported containing the number of detections in annotation measurements.

#### Experiment 3. Validation of the role of the activin pathway in LID

### Experimental design

To verify the functional role of activin in LID indicated by our snRNA-seq analysis, we designed a LID development experiment in the presence and absence of the activin receptor ALK4/TGFß1R inhibitor, SB431542. Briefly, a separate group of mice underwent unilateral 6-OHDA MFB lesions, was assessed for vehicle-induced rotations 3 weeks later as in Experiment 1 and separated into two groups: Vehicle+L-DOPA or SB431542+L-DOPA. Mice received either Vehicle (20% DMSO in 0.9% NaCl) or SB431542 (4.2 mg/kg, ip) 15 min prior to injection with L-DOPA (2 mg/kg + benserazide, 15 mg/kg, sc). A lower dose of L-DOPA was utilized compared to that used for the snRNA-seq experiments to allow for detection of either an improvement or worsening of LID due to ALK4/TGFß1R inhibition. On days 1, 3, 5, and 10 of daily treatment, AIMs and rotations were assessed as in Experiment 1. On day 11, all animals were killed 1 h after their respective treatments according to group assignment by isoflurane overdose and transcardial perfusion with 0.1 M PBS followed by 4% paraformaldehyde. Brains were post-fixed overnight in 4% paraformaldehyde and then cryoprotected with 30% sucrose in 0.1 M PBS.

### Immunohistochemistry

Cryoprotected brains were sliced on a freezing sliding microtome (40 micron) and stored in 50% glycerol/0.1 M PBS. Every 6^th^ slice containing striatum was washed in TBS before blocking for 2 h with 5% normal donkey serum. To indicate on-target efficacy of the ALK4/TGFßR1 inhibitor, primary antibodies for phospho-SMAD2 (Cell Signaling Technology, Danvers, MA, USA) and NeuN (to indicate neuronal involvement, Invitrogen-ThermoFisherScientific) were exposed overnight at 4°C to free-floating sections at concentrations (1:500-1:1000) and incubated with corresponding fluorescence-conjugated secondary antibodies (1:1000; Invitrogen-ThermoFisherScientific). Sections were also treated with primary antibodies for Iba1 (to indicate microglial involvement, Invitrogen-ThermoFisherScientific). Sections were coverslipped using Prolong Antifade Gold and stored at 4°C. Images were captured using a Nikon Ti2-C2 confocal microscope at 20X objective. Images were analyzed using QuPath as described for FISH. Nigral sections were utilized as in Experiment 1 to verify unilateral DA cell loss.

### Statistical analyses

Behavior obtained in Experiment 1 and 2 was analyzed using one-way ANOVA of averaged ALO AIMs scores for the first 60 min of behavioral ratings. TH+ immunofluorescence was analyzed by one-way ANOVA. No post-hoc analyses were warranted in either measure. Behavior in Experiment 3 was analyzed using two-way ANOVA of the summed averages of ALO AIMs scores for the entire 3 h rating period with day of exposure and drug treatment as the independent variables, followed by Bonferroni post-hoc analyses. Immunofluorescence and FISH data were analyzed using two-way ANOVAs of % of total cells expressing specific mRNA/protein, followed by Bonferroni post-hoc analyses.

## Results

### L-DOPA induced stable expression of dyskinesia in 6-OHDA-lesioned mice

Vehicle injections in 6-OHDA-lesioned mice did not induce dyskinetic movements or contralateral rotations (Figure S1B-C). As is common in 6-OHDA MFB mouse models of LID, L-DOPA treatment for 1, 5, and 10 d induced dyskinesia and contralateral rotations. The behavioral manifestations were prominent at 1 day, however, they did not significantly increase over time, despite the low dose of L-DOPA (4 mg/kg). Immunofluorescence for TH+ nigral cells revealed a similar extent of loss across all groups (Figure S1C).

### Cluster analyses of transcription reliably identified cell-type heterogeneity of the dorsal striatum

After quality control and normalization, we analyzed 97,848 cells (Intact-Naive: 13,549; DA Lesioned-Naive: 14,362; DA Lesioned-Acute 1 d L-DOPA: 26,604; DA Lesioned-Subchronic 5 d L-DOPA: 28,221; DA Lesioned-Chronic 10 d L-DOPA: 15,112). Unbiased cluster analysis revealed 27 unique clusters (Figure 1B). These clusters were characterized based on their unique marker genes and comprised all major cell types, including MSNs, interneurons, progenitor cells, and glia (Figure 1C, Table S1). Maturity status within each cell population was indicated by expression of *Tnr,* an extracellular matrix glycoprotein associated with inhibition of morphological change and highly expressed after birth (Jakovcevski et al., 2013). *Pdyn+/Drd1+* and *Penk+/Drd2+* MSNs (putatively D1- and D2-MSNs, respectively) were found to segregate independently and included multiple unique clusters. Expression of *Sema5b* and *Sgk1* has been previously associated with the patch and matrix subareas of the striatum, respectively (Martin et al., 2019; Stanley et al., 2020), and readily defined transcriptionally unique subclusters of D1- and D2-MSNs in our data set. Likewise, among matrix-localized MSNs, *Cnr1* has been associated with centrolateral topographical localization, while *Crym* is more prevalent among matrix MSNs with a medial localization (Martin et al., 2019; Stanley et al., 2020). We were also able to detect disparate subclusters of MSNs with these genes enriched. Other subclusters of cells sharing MSN marker genes were further identified based on their commonalities with so-called “eccentric” MSNs (Martin et al., 2019; Stanley et al., 2020), though little is known of their function. Furthermore, some clusters of *Drd1+* and *Drd2+* MSNs were transcriptionally distinct and subclustered independently from other MSNs, but did not meet the characteristics of Ecc MSNs. These neurons were enriched in activity-dependent gene expression programs similar to those elicited in striatal neuron cultures following KCl depolarization for 1 h (Phillips et al., 2023), and were therefore labeled as Activated (Act).

### L-DOPA exposure, but not DA lesion, differentially affects cellular proportions of striatal cells

Comparison of intact and DA-lesioned vehicle-treated striata revealed minimal changes in major cell type population ratios (ie. MSNs, interneurons, glia, oligodendrocytes, progenitors; Figure 2A-B; Table S2). In L-DOPA treated striata, the proportion of nuclei isolated from oligodendrocytes and glia nearly doubled after 1 d and 5 d L-DOPA exposure, though cellular proportions approached L-DOPA-naïve levels after 10 d of L-DOPA exposure. The oligodendrocyte effect was largely driven by oligodendrocyte progenitors and immature oligodendrocyte subtypes; radial glia, astrocytes, and microglia populations were all altered by L-DOPA treatment duration (Figure 2B, Table S2). These data are likely indicative of the supportive network that is necessary for synaptic and cellular remodeling of neurons upon L-DOPA treatment that leads to LID and will require future investigation. In addition, glial activation has been indicated previously in similar LID models and suggests an immune and/or inflammatory reaction in response to L-DOPA treatment (Elabi et al., 2023; Ferrari et al., 2021; Kuter et al., 2020; Morissette et al., 2022; Nascimento et al., 2023; Pinna et al., 2021). Though the overall populations of MSNs were not affected by L-DOPA treatment, the proportion expressing marker genes consistent with activation were increased in terms of D1-MSNs, and reduced in terms of D2-MSNs, compared to L-DOPA-naïve mice (Figure 2B, Table S2).

**Figure 2.**
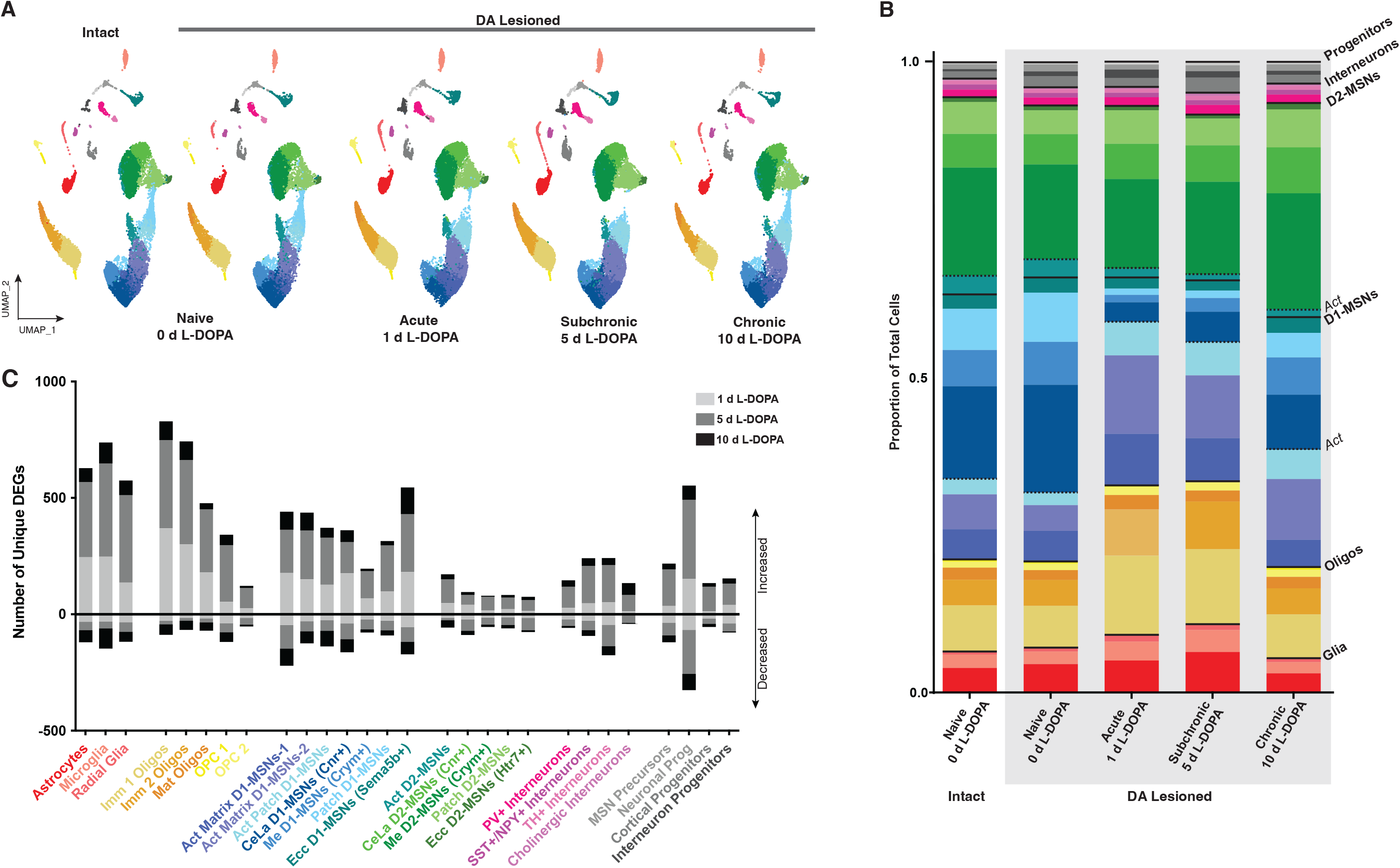
L-DOPA treatment alters major and minor subpopulation proportions and transcription within striatal cells. (A) Representative UMAPs of the striatal cellular distribution within each experimental group. (B) Bar graphs depicting the proportions of major and minor cellular subpopulations within each L-DOPA-exposure group. (C) Effect of L-DOPA exposure for 1, 5, or 10 d on the number of discrete differentially expressed genes (DEGs; log2 fold change > |0.10|) depicting either decreased or increased transcription in each striatal cell subpopulation.

### L-DOPA exposure, but not DA lesion alone, differentially affects the transcription of striatal cells

Transcriptionally, DA lesion alone resulted in <150 differentially expressed genes (DEGs) in any individual cluster (Table S4). The populations most affected DA-lesion were neural progenitor cells and subtypes of oligodendrocytes and glia; with very few changes observed in neurons. In contrast, acute L-DOPA treatment induced a vast upregulation of genes across most cellular subtypes, with the majority exhibiting >150 DEGs when comparison to L-DOPA-naïve, DA-lesioned mice (Figure 2C). Consistent with previous findings, interneurons and D2-MSNs were minimally affected by L-DOPA treatment, whereas D1-MSNs, glia, and oligodendrocytes were predominantly affected. These effects were similarly seen following 5 d of exposure to L-DOPA. By day 10 of L-DOPA treatment, while there were fewer DEGs observed overall (though still more than L-DOPA-naive, DA lesioned striata), there were roughly equivalent numbers of DEGs up-and down-regulated among individual subpopulations. Among D1-MSNs, the preponderance of upregulated DEGs was highest among Act D1-MSNs. Among non-Act matrix D1-MSNs, the medially-localized (*Crym*+) D1-MSNs expressed fewer DEGs overall than their centrolaterally-localized (*Cnr*+) counterparts.

To grasp the holistic effects of L-DOPA treatment on striatal transcription, a pseudobulk-seq analysis was completed on all striatal cell subtypes (Table S5). In general, the transcriptional differences were more pronounced acutely, with the number of unique transcripts the highest at this time point. As observed previously, IEGs were upregulated by acute L-DOPA treatment (*Fos, Fosb, Junb,* etc.). Other genes commonly associated with LID were also detected, including *Pdyn, Arc,* and *Cytb* (Dyavar et al., 2020; Smith et al., 2016; Sodersten et al., 2014). Glial-and oligodendrocyte-associated genes, such as *Cx3cr1, Ptn,* and *Mbp*, were further upregulated by acute L-DOPA treatment consistent with their increased cellular proportions following L-DOPA exposure. Pseudobulk analysis of repeated L-DOPA-treated striata indicated that a substantial reduction in the number of DEGs compared to acutely L-DOPA-treated striata, although *Fos* and *Pdyn* transcription remained elevated.

*Distinct subpopulations of D1-MSNs are altered by L-DOPA treatment and undergo subtype-and exposure-dependent transcriptional changes indicative of cellular activation and remodeling*.

Though overall proportions of D1- and D2-MSNs were not altered by DA lesion or L-DOPA treatment, individual subclusters underwent extreme rearrangement in response to L-DOPA treatment that was dependent upon the length of exposure to L-DOPA (Figure 3A-C; Table S2). L-DOPA exposure in lesioned striata, but not DA lesion alone, was associated with an increased proportion of Act D1-MSNs and decreased proportion of Act D2-MSNs (Figure 3D; Table S2). In response to L-DOPA treatment, D1-MSNs expressing markers from both patch and matrix displayed activation dependent gene expression that included enhanced IEG expression causing them to subcluster separately. These different subtypes of D1-MSNs were differentially affected in population by the number of L-DOPA exposures (Figure 3A-C). The proportion of Act Patch D1-MSNs increased due to L-DOPA experience and stayed elevated through day 10 compared to lesioned, untreated striata; non-activated Patch D1-MSNs were nearly non-existent on days 1 and 5 of exposure to L-DOPA, indicating that a large proportion of this subpopulation were identified with the IEG+ cluster (Figure 3A-B). Matrix-localized Act D1- MSNs transcriptionally diverged into two unique subclusters (Act Matrix D1-MSNs-1 and Act Matrix D1-MSNs-2). These subclusters exhibited many of the same CDGs (158 genes) but were transcriptionally distinct enough to cluster separately from one another. In general, Act Matrix D1-MSNs-1 shared more CDGs with CeLa (83 genes) and Me (23 genes) D1-MSNs, compared to Act Matrix D1-MSNs-2 (71 and 12 genes, respectively), suggesting that these populations are indicative of a spectrum of activation. Indeed, both subpopulations of Matrix Act D1-MSNs acutely increased due to L-DOPA experience, though population 2 remained elevated and population 1 ebbed to untreated levels after 10 d of L-DOPA. While the matrix D1-MSN subtypes that Act Matrix D1-MSNs likely arise from can only be speculated upon, based on ratios of populations it appeared that acute exposure to L-DOPA reduced cellular populations of Me D1-MSNs and CeLa D1-MSNs. Upon further experience with L-DOPA (5 d and 10 d), the proportion of Act Matrix D1-MSNs slightly reduced with concomitant increases in non-Act Me D1-MSNs and CeLa D1-MSNs, with a greater proportion of Me D1-MSNs returning to untreated levels by day 10. As such, it seems likely that the Matrix D1-MSNs that were in an activated state were arising predominantly from the centrolateral striatum. In contrast, Act D2-MSNs were proportionally reduced by L-DOPA treatment, regardless of the duration of exposure (Figure 3C).

**Figure 3.**
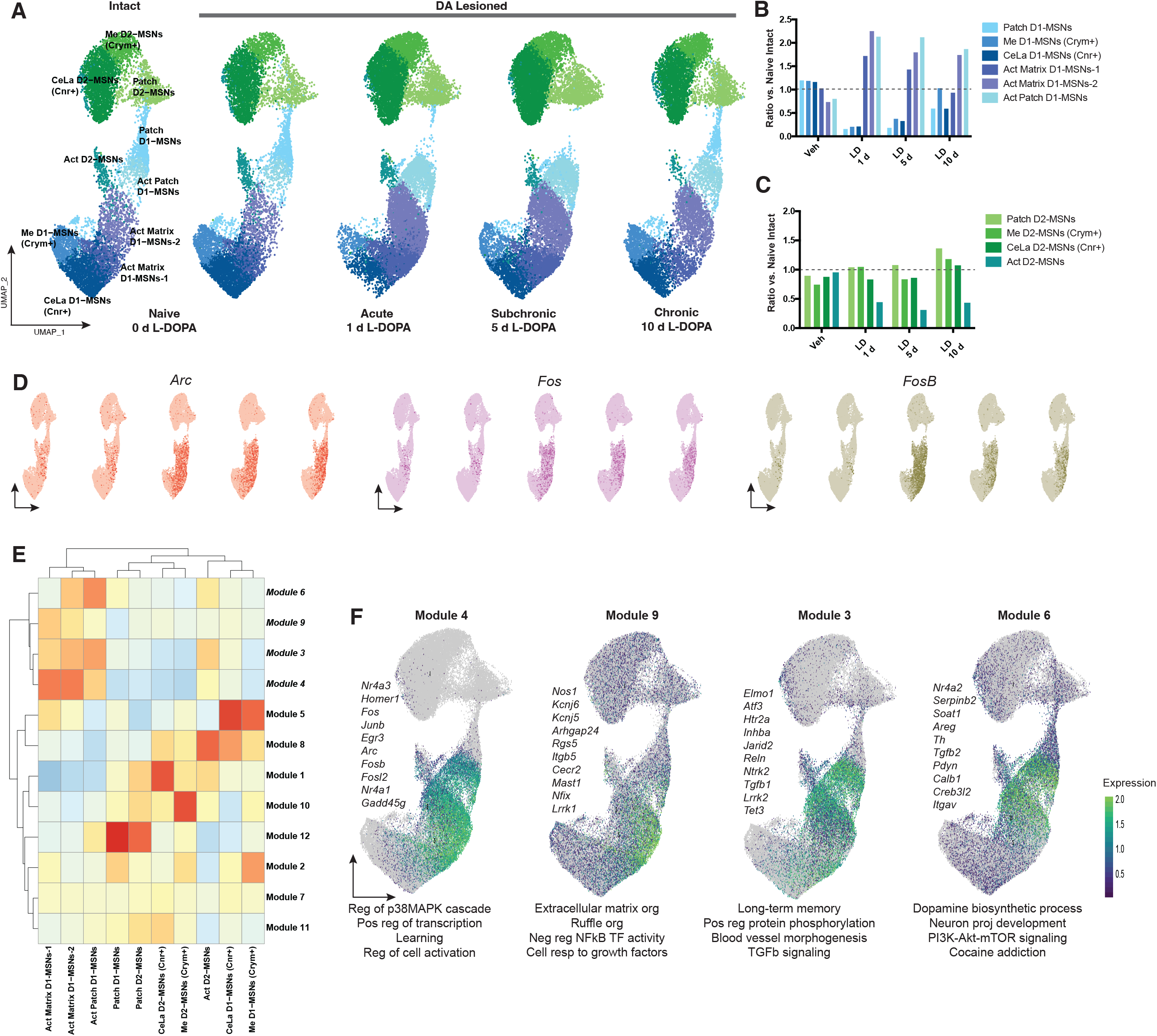
MSN subpopulations are specifically altered by L-DOPA exposure that leads to LID. (A) Representative UMAPs of the striatal MSN populations within each experimental group. (B-C) Population ratios of D1-MSN (B) and D2-MSN (C) subpopulations compared to intact, L-DOPA-naïve striata. (D) UMAPs of expression of the immediate early genes *Arc, Fos,* and *Fosb* in each experimental group. Order of UMAPs from left to right follows that depicted in *A*. (E) Gene module analysis of integrated MSNs utilizing MonocleTM. (F) Expression profiles for genes within Modules 3, 4, 6, and 9 that were enriched in Act MSNs. Selected module genes are listed next to UMAPs for each module in italics. Selected significant terms from gene ontology analysis for Modules 3, 4, 6, and 9 are denoted beneath each UMAP.

In order to better understand the transcriptional profile of each subcluster of MSN, a module analysis was completed on the integrated data (Figure 3E; Table S6). Act D1-MSNs were hierarchically distinct from all other MSN subclusters as determined by this unbiased analysis. Four modules (Modules 3, 4, 6, and 9) were also hierarchically distinct from other gene modules and were differentially enriched in Act D1-MSNs (Figure 3F). Gene ontology analyses were conducted on the genes included in each module (Figure 3F; Table S7). Act Matrix D1- MSNs-1/2 had the highest enrichment of Module 4, which included several IEGs (*Fos, Junb, Fosb,* etc.), and was enriched in gene ontology (GO) terms associated with “learning” and “positive regulation of transcription”. In comparison with Act Matrix D1-MSNs-1, Act Matrix D1- MSNs-2 expressed more genes associated with Modules 3 and 6. These modules included genes associated with “long-term memory” and “neuron projection development”. These modules were most enriched in Act Patch D1-MSNs.

To verify our results indicating the greater proportion of L-DOPA-induced transcriptional change was within D1-MSNs assigned to the activated subpopulations, we determined the enrichment of DEGs detected by Heiman et al. (2014) to our snRNA-seq data (Figure S2). Heiman and colleagues (2014) utilized a *Drd1*-TRAP methodology in 6-OHDA-lesioned, hemi-parkinsonian rats chronically treated with L-DOPA. We mapped the expression of DEGs observed within their low-dose (6 mg/kg) L-DOPA-treated group onto our integrated MSN UMAP. DEG co-expression was enriched in Act D1-MSNs, particularly Act Patch and Act Matrix-2 subclusters.

### Transcriptional effects of DA lesion and LID entrainment and reconsolidation on striatal D1- MSNs

In order to observe overall DA-lesion and L-DOPA exposure-induced changes in D1- MSNs and D2-MSNs, a pseudo-bulk analysis was completed using Monocle. Changes in gene transcription in D2-MSNs were minimal, while those in D1-MSNs were robust (Table S8). Likewise, DA lesion alone did not positively affect the transcription of many genes in MSNs (Table S8), but in D1-MSNs led to reduced expression of active transcription of genes involved in cellular activation and synaptic transmission, including glutamatergic signal transduction and G-protein coupled signaling (Table S9). In addition, genes associated with synaptic remodeling (*Reln, Ntrk, Homer1*) were also reduced by DA lesion, indicative of a loss of synaptic plasticity (Belujon et al., 2010; Picconi et al., 2003; Thiele et al., 2014).

The comparison of acute and repeated (5 and 10 d) L-DOPA exposure to vehicle treatment in D1-MSNs revealed an evolution of transcriptional activity across the development of LID (Figure 4A-C). Early in L-DOPA exposure, D1-MSNs exhibited a coordinated, bidirectional gene response indicative of enhanced DNA transcription activation and reduced protein degradation (Figure 4A2). Expression of *Drd1* was reduced by acute L-DOPA, perhaps as a compensatory response to the influx of DAergic stimulation. The upregulation of genes in D1-MSNs by acute L-DOPA was much stronger in comparison, with >120 genes with over two times greater expression in comparison to those isolated from vehicle-treated striata (Figure 4A1). This massive upregulation of genes was coupled with enhanced expression of multiple transcription factors involved in the complex formation of the pioneer transcription factor, AP-1 *(Jun, Fos, Fosb,* etc.) and transcriptional activation (*Crem, Stat6, Ell2, Smarca5, Tet3, Ebf1, Per1,* etc) (Figure 4A3). The PD risk gene, *Lrrk2* was upregulated and associated strongly with other upregulated genes associated with protein modification. TF enrichment through LISA Cistrome analysis indicated that *Creb1* was highly associated with acute L-DOPA exposure, as well as *Fos, Fosb, Mef2a* among others (Figure 4B3; Table S10).

**Figure 4.**
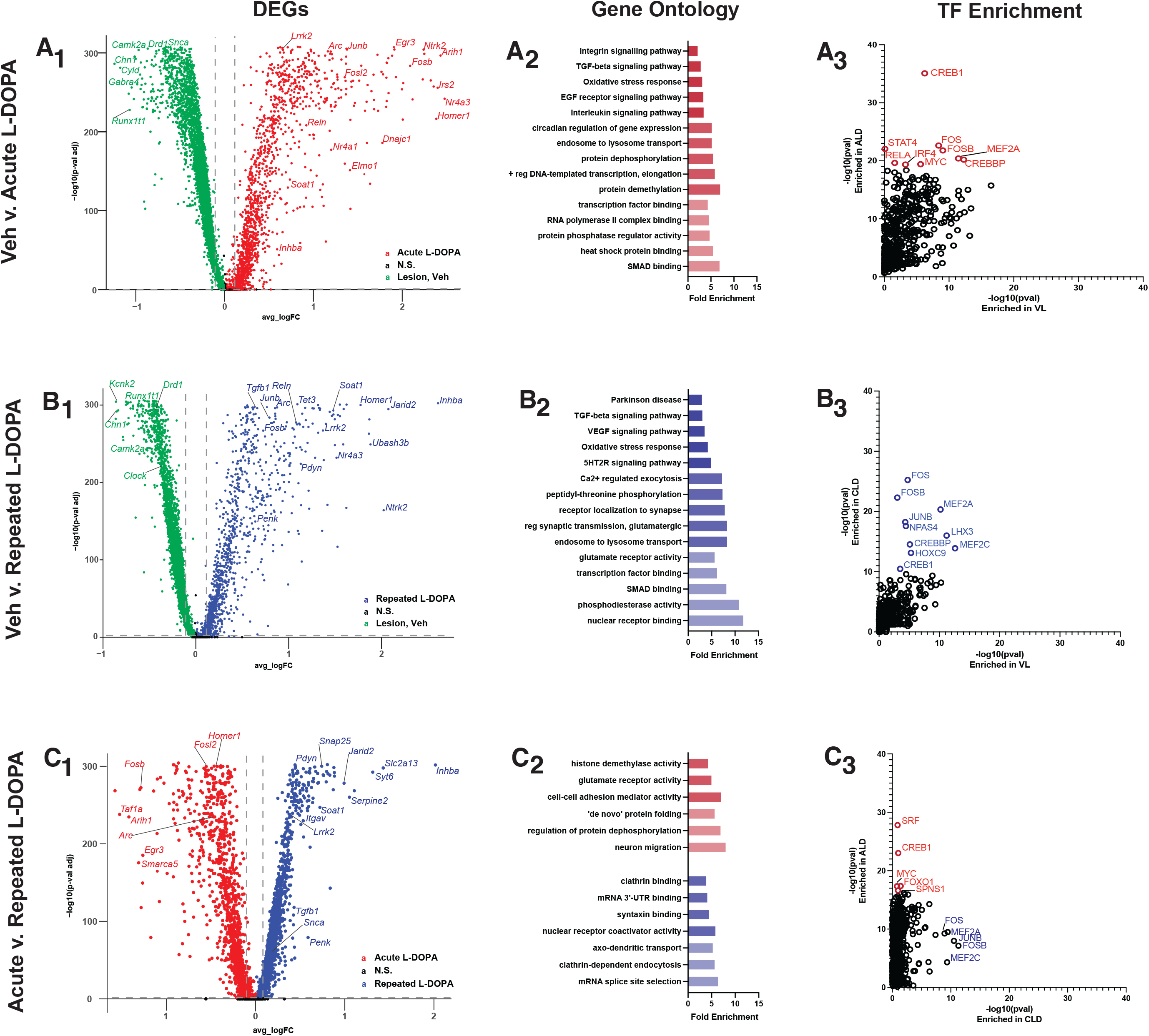
Transcriptional effects of DA lesion and LID entrainment and reconsolidation revealed by pseudobulk-seq analysis of D1-MSNs. Naïve, lesioned (green) compared to (A) acute (red; 1 d) L-DOPA-treated, lesioned or (B) repeated (blue; 5 and 10 d) L-DOPA-treated, lesioned striatal D1-MSNs. (C) Acute compared to repeated L-DOPA-treated, lesioned striatal D1-MSNs. (i) Volcano plots depicting DEGs (log2 fold change > |0.10|) with selected genes highlighted. (ii) Selected significantly enriched terms chosen from GO analyses of DEGs enriched in L-DOPA-treated groups. (iii) LISA-generated enriched TFs associated with DEGs. (D) Comparison of LID D1-MSN transcriptional profile to that observed in engram cells upon initial exposure and retrieval of a hippocampal learning paradigm in Marco et al., 2020. Analysis of DEGs upregulated upon (i) initial exposure vs retrieval, (ii) retrieval vs initial exposure, and (iii) retrieval vs basal hippocampal state in Marco et al., 2020 were compared to our striatal dataset and overlaid onto integrated striatal D1-MSN UMAPs.

Repeated L-DOPA exposure was positively associated with many of the same DEGs as revealed by acute L-DOPA, though the strength of the response (LogFC) was attenuated in most cases (Figure 4B1, Table S8). DEGs related to physical changes in neural structure and modulation of synaptic transmission were enriched after repeated L-DOPA in D1-MSNs, while those associated with regulation of membrane potential were negatively associated compared to untreated mice (Figure 4B2, Table S9). TF enrichment of *Fos, Fosb,* and *Mef2a/c* was indicated in D1-MSNs from mice undergoing repeated L-DOPA treatment (Figure 4B3; Table S10).

To better understand the processes that differentiate entrainment (first exposure) and reconsolidation (repeated exposure), the D1-MSN response to acute L-DOPA was compared to that observed in subchronic and chronic groups (Figure 4C1, Table S8). Unsurprisingly, IEGs were significantly enriched in D1-MSNs from striata of acute compared to repeated exposure groups. Other genes more strongly associated with entrainment were involved in protein dephosphorylation and folding, histone demethylase activity, and glutamate receptor activity (Figure 4C2, Table S9). In contrast, repeated exposure resulted in enrichment of genes linked to mRNA splicing and modulation of synaptic transmission. In an attempt to identify putatitive TFs responsible for the differences among acute and repeated exposure groups in D1-MSNs suggested a role for *Creb1*, *Srf*, *Myc*, and *Foxo1* in the regulation of the genes expressed following acute L-DOPA, whereas repeated exposure induced genes that were dependent upon *Mef2a*/*c*, *Fos*, *Junb*, and *Fosb* for their expression (Figure 4C3; Table S10).

### A unique role for activins within striatal D1-MSNs is indicated in the stable expression of LID

Pseudobulk analyses of D1-MSNs indicated that the gene most differentially affected by repeated exposure to L-DOPA in comparison to a single exposure was *Inhba*, which encodes for the protein inhibin subunit ßA, a member of the transforming growth factor-ß (TGFß) superfamily (Figure 4C1, Table S8). UMAP mapping (Figure 5A), as well as cluster-specific DEG analyses (Table S4, S11), revealed *Inhba* was robustly induced in patch and matrix Act D1-MSN subclusters as animals received repeated exposure to L-DOPA. To verify this result, a separate group of hemiparkinsonian mice treated acutely or chronically with L-DOPA or vehicle were utilized for localization and quantification. Analysis of striatal tissue using FISH indicated that both acute and chronic L-DOPA led to upregulation of co-expression of *Fos* transcript in *Drd1+* cells compared to untreated mice, though levels were highest in acutely treated mice (Figure 5C-D). This result was also indicated in our snRNA-seq dataset (Fig 5B). A significant induction of *Inhba* transcription in *Drd1*+ cells was observed in animals with chronic L-DOPA exposure, in comparison to acutely treated or untreated mice (Figure 5C-D). Furthermore, chronic L-DOPA induced a significant increase in *Inhba* transcript in those *Drd1+ cells* that were also *Fos+* (likely Act D1-MSNs), corroborating results obtained using snRNA-seq.

**Figure 5.**
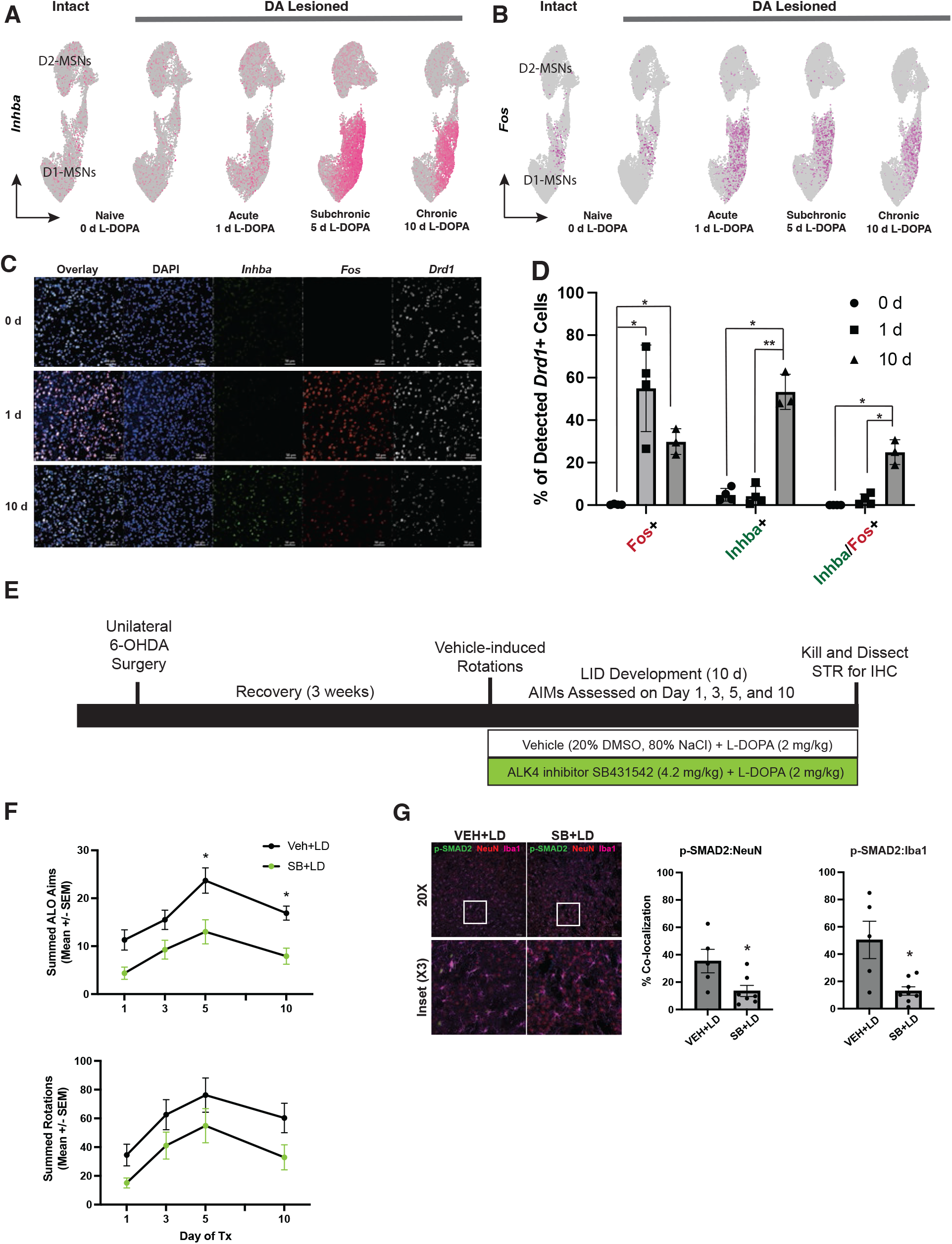
*Inhba*/Activin/ALK4 signaling is enhanced by repeated L-DOPA treatment in *Fos*+ D1- MSNs and modulates LID development. (A) UMAPs of expression of *Inhba* and *Tgfb1* mRNA in each experimental group. (B) Confocal dorsal striatal images (20X) depicting enhancement of *Inhba* transcript in *Fos+/Drd1*+ cells in animals treated for 10 d L-DOPA compared to those treated for 0 or 1 d with L-DOPA. (C) QuPath assisted quantification of confocal images depicting *Inhba, Fos,* and *Drd1* transcript co-expression in hemiparkinsonian mice treated for 0, 1, or 10 d L-DOPA. (D) Time course of experiment and graphical description of experimental groups utilized to establish the functional role of ALK4/TGFb1R signaling in LID development. (E) Effects of ALK4 inhibition with SB431542 on summed ALO AIMs (top) and contralateral rotations (bottom) observed on days 1, 3, 5, and 10 of co-treatment with L-DOPA. (F) Confocal images (20X) depicting pSMAD4 localization in NeuN+ and Iba1+ cells in the dorsal striatum of hemiparkinsonian mice treated with SB421542 (0 or 4.2 mg/kg) and L-DOPA for 10 d and QuPath assisted quantification of confocal images depicting pSMAD4, NeuN, and Iba1 co-expression in mice treated with SB421542 (0 or 4.2 mg/kg) and L-DOPA for 10 d.

### *ALK4R/TGF*ß*R1 inhibition reduces development of LID*

*Inhba* encodes for a preproprotein that is cleaved to form ßA subunits of activin and inhibin protein complexes (Bloise et al., 2019). Dimerization of the ßA subunit to another ßA subunit results in the formation of the Activin A protein complex, while dimerization with an α subunit results in formation of an inhibin B complex. Since transcription of other subunits of inhibin were not significantly affected by L-DOPA in D1-MSNs, we suspected that formation of Activin A (ßA / ßA) protein complexes would be most likely to occur. Activin A has been shown to modulate cellular differentiation, immune function, and neuroplasticity (Ageta et al., 2010; Bloise et al., 2019; Gancarz et al., 2015). To explore the possibility of a functional role of the activin/TGFß pathway in LID as indicated by our snRNA-seq analysis and corroborated by FISH, we designed a LID development experiment in the presence and absence of the ALK4R/TGFßR1 antagonist, SB431542 (Figure 5E; (Inman et al., 2002)). At doses previously utilized to influence hippocampal synaptic plasticity and did not interfere with overall motor activity (Caraci et al., 2015), SB431542 dampened development of LID over the course of 10 d of treatment with L-DOPA, compared to vehicle (Figure 5F; F_Drug_(1,19)=13.43, p=0.0016). This effect reached statistical significance by d 5 and continued to d 10 (both p<0.01), suggesting that reducing activin/TGFß signaling can inhibit the stable expression of LID. On target efficacy was verified by reduced pSMAD nuclear localization in both neurons and microglia (Figure 5G), which have the highest expression of TGFßR1 in the brain (Table S3), in hemiparkinsonian mice which received concomitant SB431542 and L-DOPA in comparison to L-DOPA alone.

## Discussion

Here we report a comprehensive profile of the striatal transcriptional landscape following the acute and chronic exposure to L-DOPA in dopamine-depleted animals, identifying specific subpopulations of D1-MSNs that are uniquely involved in LID development and maintenance. We corroborate prior sn and scRNA-seq investigations into striatal cellular heterogeneity (Gokce et al., 2016; Martin et al., 2019; Stanley et al., 2020), observing over 25 unique subpopulations, comprising both classes of MSNs, all major interneuron subtypes, progenitor cells, and glia (Figure 1B-C). While DA cell loss alone induced minor transcriptional changes in this neurotoxin model, L-DOPA exposure had wide-ranging effects on almost all observed cell populations (Figure 2). As expected, D1-MSNs were particularly sensitive to L-DOPA treatment, forming new states within both patch and matrix-residing D1-MSNs that expressed transcriptional profiles consistent with cellular activation (Figure 3). The number of cells activated as well as the transcriptional behavior of both patch and matrix D1-MSNs was directly related to the duration of L-DOPA treatment (Figure 4). In D1-MSNs, the acute exposure to L-DOPA induced several transcription factors associated with neuronal plasticity and direct regulators of MAPK signaling, while the transcriptional behavior following chronic L-DOPA was more strongly related to synaptic stabilization and TGFß signaling. Interestingly, we found a high degree of similarity between the transcriptional response of D1-MSNs to L-DOPA and the gene expression profile previously seen in hippocampal engram cells during the consolidation and retrieval of an associative memory ((Marco et al., 2020); Figure 4D). We then illustrated the translational applicability of these results, verifying that a subset of D1-MSNs shows an enhancement in Activin (*Inhba*) by FISH following chronic L-DOPA exposure, and that the pharmacological inhibition of Activin/TGFß signaling decreased dyskinesia severity (Figure 5). Collectively, these data illustrate the power of this comprehensive sequencing dataset for both hypothesis generation and validation in LID and highlight a previously unappreciated role for Activin/TGFß signaling in the development of LID.

The strength of this dataset lies in the comprehensive sequencing of the large number of cells sampled (>90,000), enabling the identification of various subpopulations of glial and neuronal cells previously challenging to distinguish (Figure 1). Consistent with previous results (Heiman et al., 2014), the loss of DA neurons had minimal effects on gene transcription, while L-DOPA exposure led to the emergence of three unique activation-associated subclusters, referred to here as Act Matrix D1-MSNs-1, Act Matrix D1-MSNs-2, and Act Patch D1-MSNs (Figure 3). While all three clusters were predominantly in L-DOPA-treated animals, each activation state displayed a unique transcriptional profile that became apparent depending upon the duration of L-DOPA treatment. While acute L-DOPA exposure caused the subclustering of D1-MSNs into all 3 activation subtypes, following repeated exposures the number of cells in the Act Matrix D1-MSNs-1 cluster began to taper off, while Act Matrix D1-MSNs-2 and Act Patch D1-MSNs remain relatively stable across 1, 5, and 10 d of L-DOPA.Our data suggests that the quantity and quality of activated neurons changes following repeated treatment, with a pruning of weakly stimulated neurons recruited in the initial circuit as well as a shift in the transcriptional behavior of the entrained neurons as the circuit matures. As such, Act Matrix D1-MSNs-1 likely represents a population undergoing the first wave of activation-induced transcription, whereas Act Matrix D1-MSNs-2 may represent a more accelerated transcriptional response, as the second wave of transcription is induced by first wave factors (Alberini, 2009). Transcriptionally, patch and matrix MSNs exhibited very similar gene expression that was associated with activation, altered cell signaling, and long-term synaptic potentiation upon L-DOPA treatment. However, our results suggest that chronic L-DOPA treatment had an outsized effect on the activation state of patch D1-MSNs compared to matrix D1-MSNs, data consistent with previous research suggesting that patch-localized MSNs play a more significant role in LID expression (Cenci et al., 1999; Crittenden et al., 2009; Henry et al., 2003; Mahmoudi et al., 2009; Saka et al., 1999).

Acute L-DOPA exposure induces a widespread cellular activation that, following repeated exposure, becomes focused to a select population of neurons (Engeln et al., 2016; Fieblinger et al., 2018; Girasole et al., 2018), processes akin to the early circuit refinement underlying learning and memory formation (Liu et al., 2012; Ramirez et al., 2013). Hippocampal-dependent memory relies upon unique ensembles of neurons known as “engrams”, that are recurrently activated during the initial formation and subsequent retrieval of a memory. An essential characteristic of engrams is that activation of this neuronal population is sufficient to drive conditioned behavioral responses even in the absence triggering stimuli (Josselyn & Tonegawa, 2020). Recent research in LID has revealed that a subset of D1-MSNs is recurrently activated during the initial and subsequent L-DOPA exposures, and the inhibition and/or excitation of these cells can bidirectionally modulate the behavioral manifestation of LID consistent with putative “engram” cells (Engeln et al., 2016; Fieblinger et al., 2018; Girasole et al., 2018). Our collective data suggest that the two persistent D1-MSN activated subpopulations (Act Patch D1-MSNs and Act Matrix D1-MSNs-2) likely represent these "engram"-like cells, storing environmental exposures in a shared motor-circuit, and dependent upon mechanisms similar to those seen in other models of synaptic plasticity including learning and memory, drug addiction, and physiologic motor learning paradigms (Josselyn & Tonegawa, 2020). In support, we compared our data set to hippocampal nRNA-seq data from an *Arc*-TRAP mouse model in which *Arc*+ cells are persistently fluorescently tagged during entrainment of a hippocampal dependent task, putative “engram” cells (Marco et al., 2020). Interestingly, the hippocampal engram cell transcriptional profile was most enriched among the more stable Act D1-MSNs (Act Patch and Act Matrix-2 D1-MSNs), as would be expected if these subpopulations are indicative of an engrammatic population that are involved in the encoding and retrieval of the dyskinesia-inducing motor response to L-DOPA (Figure S3) and strongly suggests there are shared mechanisms underlying neuronal synaptic plasticity and the storage of long-term behaviors.

While many of the differentially expressed genes (DEGs) identified following acute L-DOPA exposure were also found after prolonged exposure (Figure 4A-B), our analysis revealed several important transcripts uniquely expressed following chronic treatment (Figure 4C). Initially, L-DOPA exposure led to the induction of several genes found to define the Act Matrix D1-MSNs-1 subpopulation including multiple transcription factors necessary for synaptic plasticity, such as AP-1 and Egr family members, along with several regulators of MAPK signaling (Figure 3E). Conversely, repeated L-DOPA exposure predominantly induced two unique subpopulations, Act Matrix D1-MSNs-2 and Act Patch D1-MSNs (Figure 3B), that were related to synaptic remodeling and transmission, and included genes such as *Ntrk2*, *Elmo1*, and *Soat1* ((Kim et al., 2011; Musumeci et al., 2009); Figure 3E). Underlying these temporal differences in transcriptional behavior were distinct transcription factors, with acute L-DOPA administration causing Creb1, Srf, Myc, and Foxo1 to induce the expression of IEGs, such as *Fos*, *Jun*, and *Egr1*. However, after 5-10 d of L-DOPA treatment, the transcriptional network became increasingly dependent upon the previously inducible factors: Mef2a/c, Fos, Junb, and Fosb. This transition away from calcium-dependent factors for the underlying transcriptional regulation strongly resembles the patterns found in alternative models of long-term behavioral memory, where many of these same inducible factors play essential roles in establishing the necessary transcriptional network for multiple models of hippocampal and striatal plasticity (Alberini & Kandel, 2014; Cenci, 2002). In sum, these data indicate that early exposures to L-DOPA are inducing transcriptional activity to initiate synaptic remodeling that is dependent upon canonical neuronal signaling; however, as the behavioral sensitization increases in concert with the synaptic strength, the transcriptional network becomes increasingly dependent upon the sustained expression of these previously temporarily inducible factors.

Based upon our pseudobulk RNA-seq analysis of D1-MSNs, one of the genes specifically associated with chronic L-DOPA treatment was *Inhba*, which encodes inhibin subunit βA, a member of the TGFβ superfamily that acts as a ligand for ALK receptors. In D1- MSNs, *Inhba* expression was over 4 times higher in mice repeatedly treated with L-DOPA compared to those with acute exposure, suggesting that it may be involved in behavioral sensitization to L-DOPA. Blocking ALK/TGFβ receptor signaling using SB431542 attenuated LID development, suggesting an essential role for Activin in LID development. While some version of neuronal autocrine signaling is possible, *Inhba* expression was also elevated in several glial populations, including oligodendrocytes and microglia (Table S3). Activin has been implicated in microglial activation and oligodendrocyte maturation (Dillenburg et al., 2018; Morianos et al., 2019; Wang et al., 2022); however, very little is known about the importance of glial or neuronal Activin/TGF-β signaling in striatal plasticity and future work identifying the role each cell type plays in this signaling pathway will be essential for therapeutic development.

Collectively, these data provide a comprehensive view of the transcriptional changes that underlie the aberrant striatal plasticity induced by L-DOPA that causes LID, while also providing several promising avenues for future research. Due to the extensive transcriptional changes that correlate to LID development, in this study we have focused on the disparate activation states of D1-MSNs induced by L-DOPA treatment, showing numerous parallels between the mechanisms necessary for associative learning and those underlying LID. These data strongly suggest that maladaptive synaptic plasticity directly contributes to the progressive L-DOPA sensitivity, and that the neurons activated during initial L-DOPA exposure are molecularly “primed” so that subsequent L-DOPA treatments unleash an imprinted motor program stored in these putative “engram cells” causing dyskinetic behaviors. The targeting of these individual cells may hold promise for inhibiting the development of LID or extinguishing a previously formed cellular “memory” for L-DOPA, perhaps by targeting activin-mediated mechanisms. These data represent the first step toward establishing and delineating the unique roles of individual subsets of striatal cells in LID and start to define the differential gene modules necessary for the development of the aberrant circuit caused by dysregulated dopaminergic signaling.

## Acknowledgements

This work was supported by grants from the American Parkinson Disease Association (APDA) and the Parkinson Association of Alabama to Dr. Jaunarajs as well as the APDA Advanced Center for Parkinson Research at UAB. We sincerely thank the UAB Flow Cytometry and Single-Cell Core Facility, particularly Drs. Shanrun Liu and Vidya Sagar Hanumanthu, for their assistance in protocol development, and Drs. Jeremy Day (UAB-Comprehensive Neuroscience Center) and Ashley Harms (UAB-Center for Neurodegeneration and Experimental Therapeutics) for their guidance in method development and interpretation.

## Supplemental Files

Table S1-11. Contains data for snRNA-seq analyses, including cluster defining genes, DEG lists, TF enrichments, Monocle gene module analyses, and GO analyses. Descriptions for each table are described on the first line of each table.

**Figure S1.**
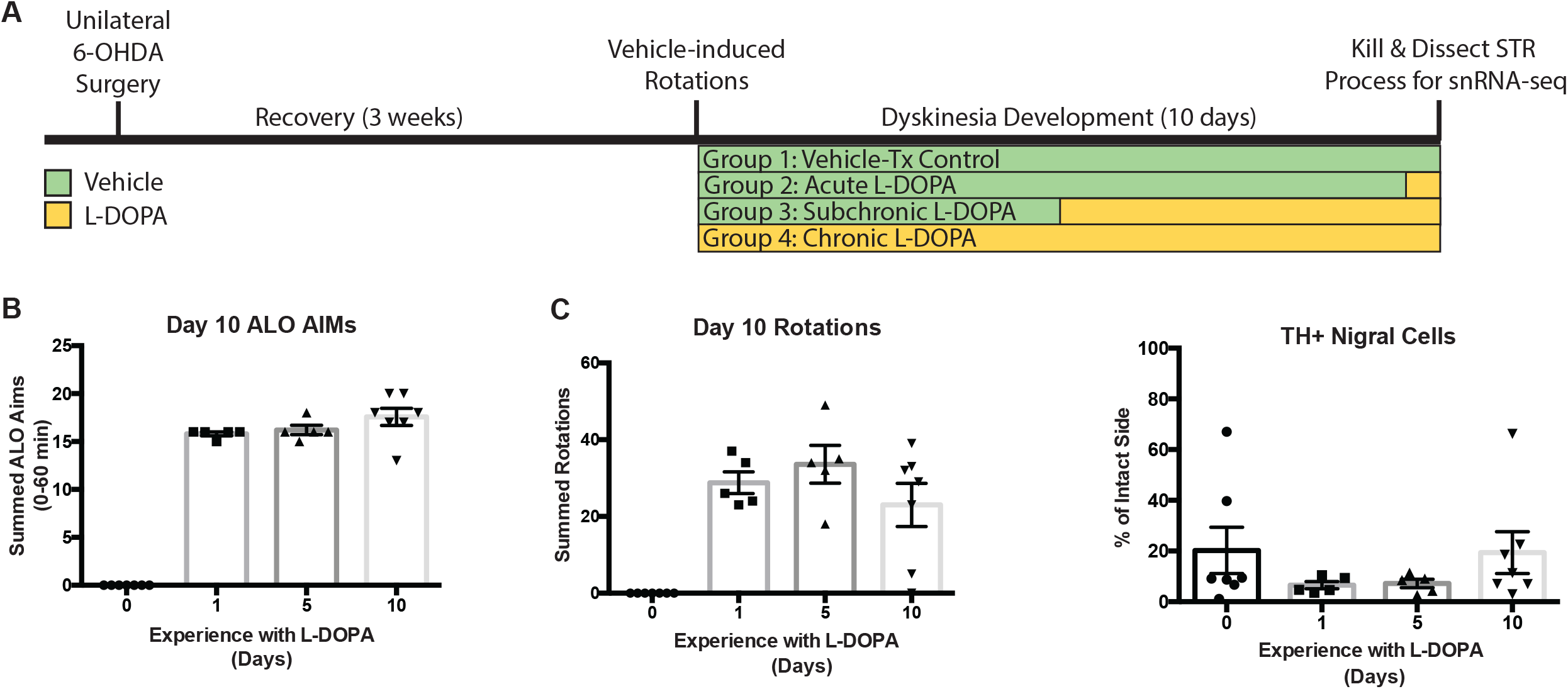
Behavior and dopamine-depletion data for Experiment 1. A) Time course of Experiment 1 experimental design. B) ALO AIMs and C) contralateral rotations (+/- SEM) for the first 60 min of the final behavioral session for each group of animals before tissue harvesting. D) Percentage of TH+ cells in nigral sections on intact and DA lesioned side (+/- SEM).

**Figure S2.**
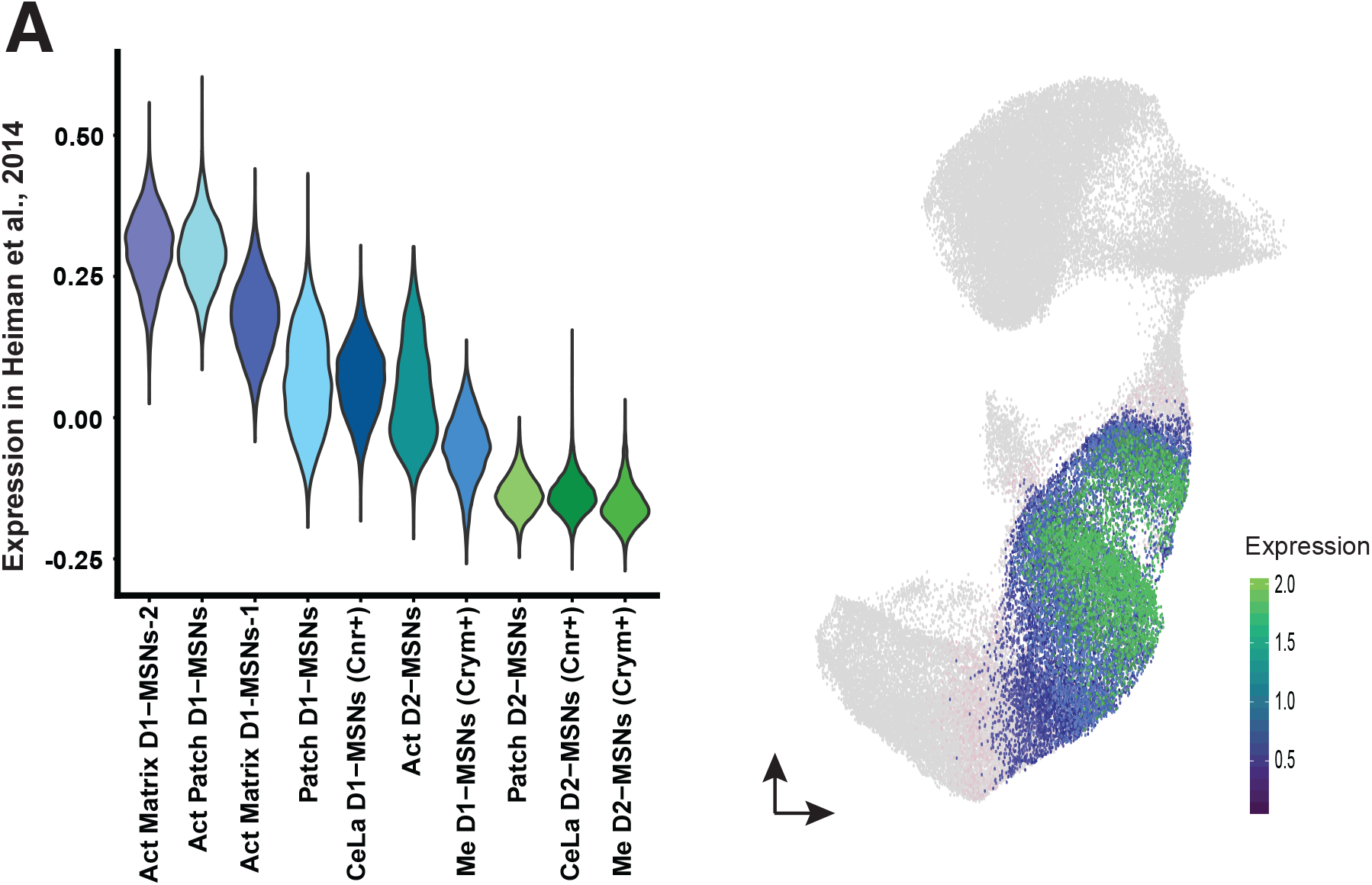
Expression profile of DEGs observed in RNA-seq derived from a Drd1a-BAC-TRAP model of LID (Heiman et al., 2014) mapped onto MSNs from the current snRNA-seq dataset. (A) Violin plot depicting ratio of co-expressed DEGs within integrated MSN subpopulations and (B) mapped enrichment of the current dataset compared to that published by Heiman et al., 2014.

**Figure S3.**
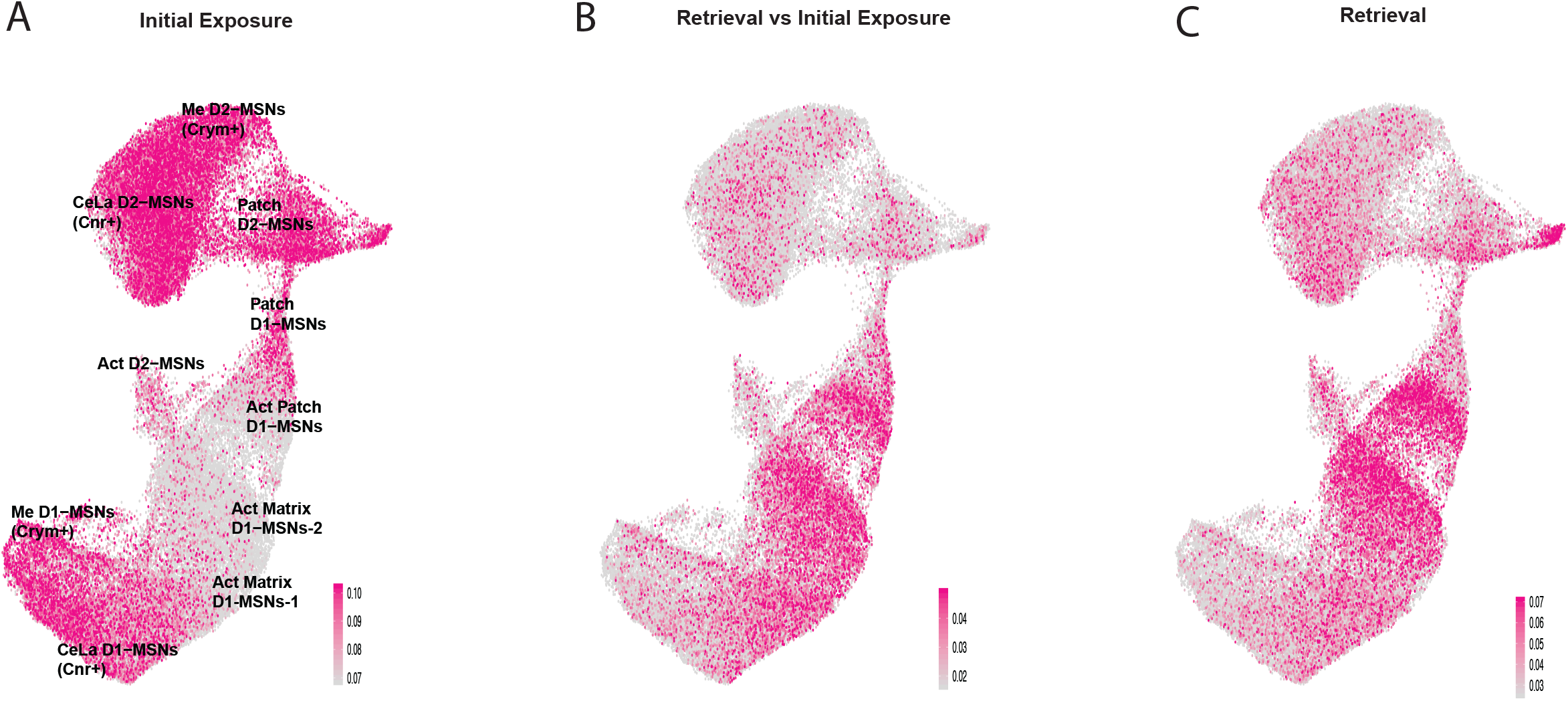
Expression profile of DEGs observed in nRNA-seq derived from an Arc-BAC-TRAP model of associative learning (Marco et al., 2020) mapped onto MSNs from the current snRNA-seq dataset. (A-C) Mapped enrichment of DEGs from the current dataset compared to those observed after the (A) initial exposure to the stimulus compared to homecage controls, (B) the retrieval of the stimulus-response association compared to the initial exposure, and (C) the retrieval of the stimulus-response association compared to homecage in Marco et al., 2020.

## Notes

### Competing Interest Statement

The authors have declared no competing interest.

### Summary of Updates

Improved figure quality.

